# A proteome-based design of bitter peptide digestion regime to attenuate bone soup bitterness: comparison with a rainbow trout extract-mediated bitter taste masking approach

**DOI:** 10.1101/279265

**Authors:** Ying Han, Changlu Guo, Zhengyu Yan, Feng Jin, Jie Jiang, Zhizhou Zhang

## Abstract

**BACKGROUND:** The fresh bones (with some meat on them; frequently discarded as a large quantity of industry garbage) of marine fish such as cod and salmon are good materials for manufacture of food additives (taste adjusters). However, such fish-bone originated additives often have apparent bitter taste and need additional debittering regime.

**RESULTS:** In this study, 46 known bitter peptides in the cod proteome were targeted for specific protease digestion to eliminate bitter taste from the cod bone soup. Though the debittering effect was apparent, the bitter taste was not completely removed. However, the bitter taste can be removed by addition of trout extract to a complete extent. The strong debittering power of rainbow trout extract was further confirmed by the debittering experiments on salmon bone soup and bitter melon, both with perfect results.

**CONCLUSION:** These results indicated that the cod bone soup bitterness comes not only from bitter peptide but also from other substances that can be masked by trout extract. Considering the fact that trout proteome has more potential bitter peptides than cod, trout extract shall have a strong bitter masking substance to be determined in the future.

## INTRODUCTION

Many kinds of food will form a bitter taste after the hydrolysis, seriously affecting the market prospects. The bitterness of a protein hydrolates is largely due to the release of bitter peptides, while many bitter substances other than bitter peptides have not yet been well identified. Common bitterness in a food is often derived from cholic acid or bitter amino acids (histidine, leucine, isoleucine, methionine, etc.). Fish tissue waste (skin, tail, bones, fishbone, fins, visceral, etc.) is often mixed with bile or bile acid and other bitter substances, resulting in a taste of obvious bitterness. In principle, it is possible to eliminate the cholic acid or bitter amino acids through specific biotransformation reactions. At present, there are at least 46 known bitter peptides^1-10^ (Table 1), but only 15 bitter peptides were listed in the BitterDB database.^11^

**Table 1.**
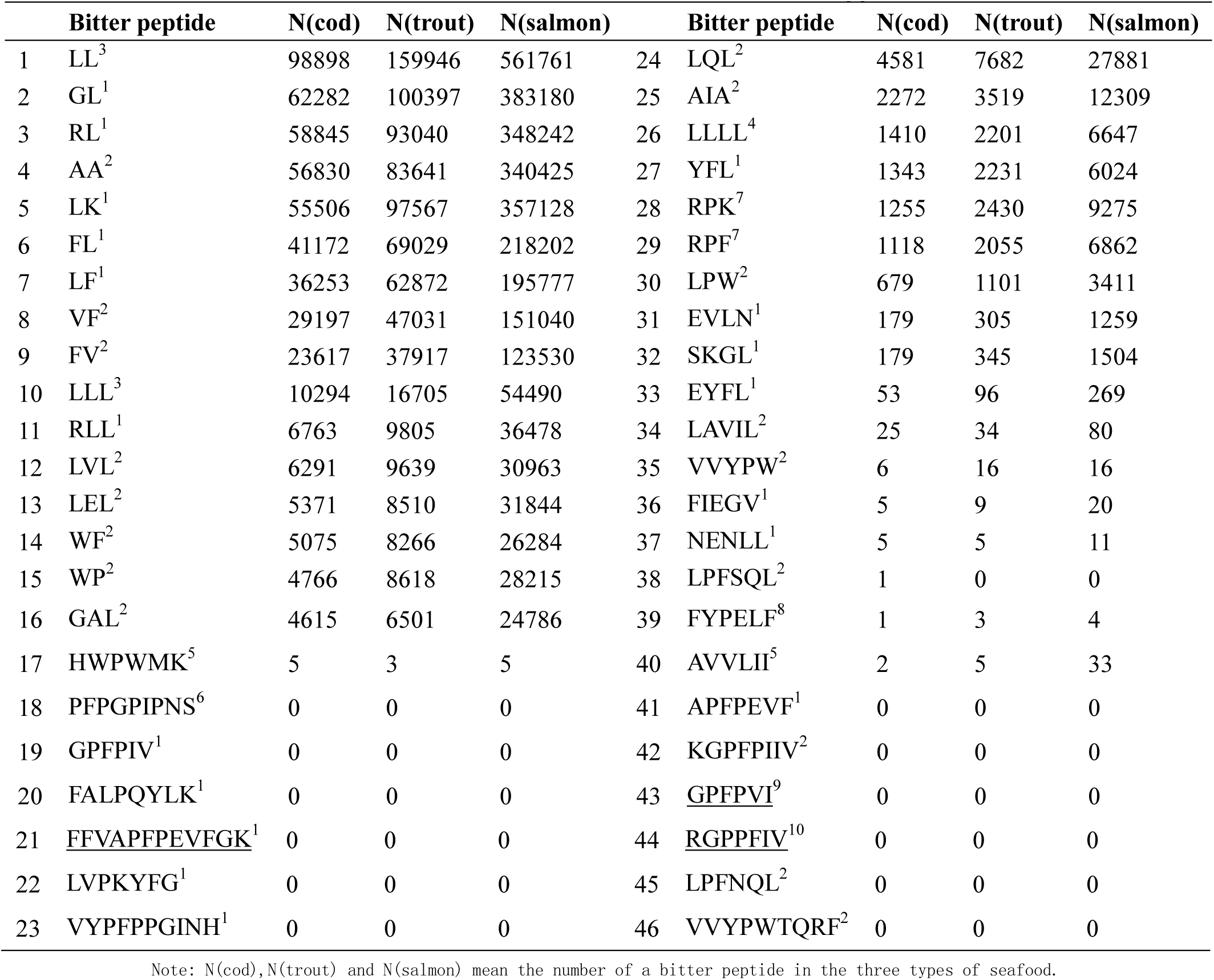
Predicted numbers of 46 bitter peptides in three types of seafood

Various protein hydrolysates with bitter tastes were found from 1950s.^12^ Since then, researchers have proposed debittering methods such as activated-carbon adsorption, chloroform extraction, ethanol extraction, isoelectric point precipitation, hydrophobic interaction chromatography, and bitter masking.^13-18^ Some types of food are hard to de-bitter and need combinations of different technologies and approaches for satisfactory debittering effect, but possibly resulting in high cost of food processing.

Few reports have been seen in that the whole proteome sequences are directly employed to quantify the types of potential bitter peptides and thereby design specific enzyme cutting strategy for those potential bitter peptides. In this study, bitter peptide breakage regimes based on cod proteome sequences were set up, but it was found that such regimes were not able to well eliminate the bitter taste of cod bone soup. Trout bone soup didn’t taste bitter at all though its proteome has more potential bitter peptides than cod. Salmon proteome has more potential bitter peptides than both cod and trout, but its bone soup was similarly as bitter as cod bone soup. Besides, mixing trout bone soup with either cod bone soup or salmon bone soup completely eliminated the bitter taste, suggesting that trout bone soup has some bitter-masking substance. Surprisingly, even the bitter taste of bitter melon^19^ soup was also mostly or completely eliminated by trout bone soup.

## MATERIALS AND METHODS

### Bone soup

All kinds of fresh fish bone were provided by The YueYi Biotech Ltd, RiZhao, Shandong, China. Cod fish bone was equally mixed with water (W/W), smashed with a stirrer to get a cod bone suspension. The suspension was equally mixed with water (V/V) and then heated to the boiling state for 20 min to get the cod bone soup. Other kind of bone coup mentioned in this study was made the same way. Amino acids and fatty acids compositions in three types of bone soup were measured (averaged from three repetitions) in Merieux Nutrisciences, Sino Analytical (Qingdao) Ltd, China.

### Determination of potential bitter peptide contents in a proteome

In the database BitterDB (http://bitterdb.agri.huji.ac.il/dbbitter.php#Home)^11^, there are 680 bitter compounds and only15 bitter peptides included. In this study, total 46 bitter peptides (Table 1) were included (Some other bitter peptides are not included in this study at the time of submission). They were scanned and calculated along the whole proteome using a laboratory software program designed by the authors or simply search and calculate using the office software WORD’s basic function. Proteome sequences of Cod (Gadus morhua) were obtained from http://www.ensembl.org/Gadus_morhua/Info/Index; Proteome sequences of rainbow trout (Oncorhynchus mykiss) were downloaded from https://www.ncbi.nlm.nih.gov/genome/?term=Oncorhynchus+mykiss; and the proteome sequences of Salmosalar (Atlantic salmon) were obtained from https://www.ncbi.nlm.nih.gov/genome/369. Because proteins in the fish bone belong to part of the whole proteome, bitter peptides scanned from the whole proteome shall cover those in the fish bone soup.

### Proteases used to digest the whole proteome

The enzymatic specificity characteristics of several proteases were listed in Table 2. In order to theoretically cut apart all twenty kinds of amino acids, several combinations of proteases can be considered as follow. (1) Carboxypeptidase A, B and C can work together to cut all amino acids; (2) Bromelain and protease K are another potential combination; (3) Pepsin, trypsin and Bromelain can work as a full functional combination; (4) Pepsin, trypsin and Carboxypeptidase A are also a good combination; Apparently there are also several other different combinations capable of digesting all twenty amino acids. However, cost efficiency is an important consideration in practice. Protease K and Carboxypeptidases are much more expensive than normal proteases such as pepsin, trypsin and Bromelain. So the combination of Pepsin, trypsin and Bromelain was selected in this study. Pepsin (Cat.Num.150624, 12k U g^-1^), Trypsin (Cat.Num.150726, titer 1:250), and Bromelain (Cat. Num.150618, 800k U g^-1^) were purchased from Beijing HongRunBaoShun Scientific Ltd. Company, China. The general regime included (1) all cuttings were set carboxyl-terminal; (2) R and K residues were subjected to Trypsin; F, Y, W, D, E, A, V, L, I, P, M residues were subjected to pepsin; and (3) all other residues were in charged by Bromelain.

**Table 2.**
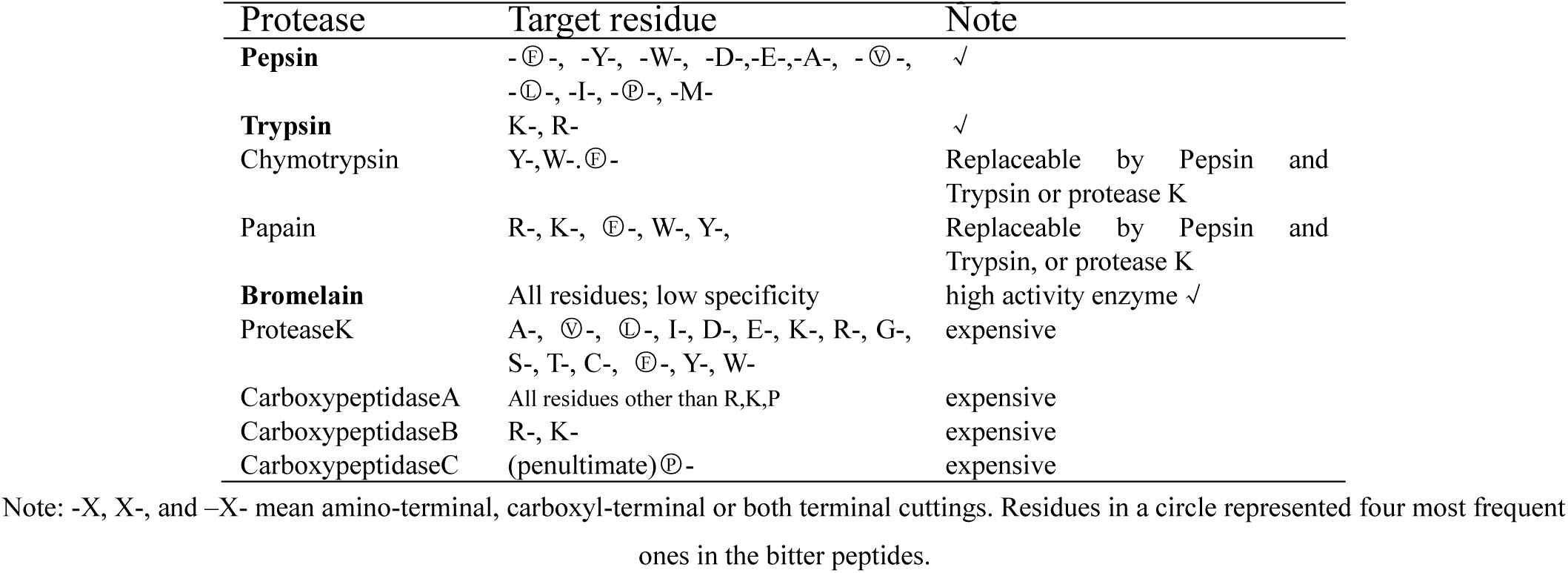
Proteases used to cut peptide bonds

### Proteases used to cut the bitter peptides

The above protease combination was also applicable for bitter peptide digestion away from the proteome. The most frequent residues in a bitter peptide were L, P, F and V, and less frequent bitter-peptide residues were G, H, A, Y, R, E and R. So the pepsin-trypsin-bromelain combination shall be good for bitter peptide digestion. For any bitter peptide –NNNX_1_X_2…_X_n-1_X_n_NNN (X, bitter peptide residue; N, neighboring non-bitter peptide residue), a general digestion regime was undertaken as –NNNX_1_↓X_2_↓_…_X_n-1_↓ X_n_ NNN, in which the cutting arrows were carboxyl for X_1_to X_n-1_ respectively.

### Sensory evaluation

The bitterness of each soup sample was estimated (averaged with three repetitions) by the quinine-sulfate (purchased from Sangon, Shanghai) equivalent test. The bitterness of a sample was compared with a series of quinine-sulfate dilutions using a sensory panel composed of five people. The panelists were trained with standard quinine-sulfate solutions set at several concentrations near the threshold levels. The panelists rinsed their mouths thoroughly with water and then keep 2ml soup in the mouth for 10 sec before evaluation.The degree of bitterness was rated as not (-), slightly (+), distinctly (++), moderately (+++), very (++++) and extremely bitter (+++++). Each degree represented a quinine-sulfate concentration of 1.6, 2.4, 3.2, 4.0, 4.8 and 5.6×10^-5^ mol L^-1^, respectively.

## RESULTS

### Bitter peptide profiles in cod, salmon and rainbow trout

According to EBI genomes and the relevant specialized website for complete proteomics information, the contents of bitter peptides of the three species were obtained as follows (Table 3). Cod, salmon and trout have different sizes of genomes, so their proteomes. This is the main reason why they had different numbers of bitter peptides (Table 1). Some longer bitter peptides include several shorter bitter peptides in them, so cutting of longer bitter peptides will release shorter ones that might be more bitter. So complete enzymatic cutting was expected in order to finally reach individual amino acid residues.

**Table 3.** Enzymatic cutting site numbers of each protease in three proteomes

| Protease | cod | salmon | trout | cod | salmon | trout |
| --- | --- | --- | --- | --- | --- | --- |
|  | Bitter peptide cutting |  |  | Whole proteome cutting |  |  |
| Pepsin | 427259 | 2438483 | 694866 | 5562656 | 39200011 | 9635931 |
| (amount) | (6g) | (34g) | (9.8g) | (78g) | (550g) | (135g) |
| Trypsin | 68160 | 402361 | 107675 | 1127717 | 8155897 | 1987439 |
| (amount) | (0.2g) | (1.2g) | (0.3g) | (3.3g) | (24g) | (5.8g) |
| Bromelain | 71858 | 411021 | 115282 | 3329959 | 25010690 | 6188981 |
| (amount) | (12mg) | (69mg) | (19mg) | (556mg) | (4.2g) | (1g) |
| Total protease mixture (g) | 6.2 | 35.2 | 10.1 | 81.8 | 578.2 | 141.8 |

### Effects of bitter peptide cutting on debittering of the bone soup

Fresh cod fish soup 600ml was heated to boil for 20min, then cooled to room temperature and divided into four equal parts (150 ml each) into the same four containers. Protease mixtures were added according to Table 4 to purposely digest bitter peptides. Bitter taste was evaluated at three time intervals. Some residual bitter taste was still sensed after 24 hours. Similar results were seen for salmon soup (data not shown). Since the trout soup was found not bitter at all, so the bitter peptide cutting test was not undertaken on it.

**Table 4.** Debittering effect of bitter peptide cutting in the cod proteome

| No. | Total protease mixture (g) | Bitterness (1hr) | Bitterness (6hr) | Bitterness (24hr) |
| --- | --- | --- | --- | --- |
| 1 | 0 | ++++ | ++++ | ++++ |
| 2 | 0.62 | ++++ | ++ | + |
| 3 | 1.24 | +++ | + | + |
| 4 | 6.2 | +++ | - | + |

### Effects of whole proteome cutting on debittering of the bone soup

Similarly as above, fresh cod fish soup 600ml was heated to boil for 20min, then cooled to room temperature and divided into four equal parts (150 ml each) into the same four containers. Protease mixtures were added according to Table 5 to purposely digest the whole proteome (Table 3). After 30 hours, there was still some residual bitter taste in the samples. Similar results were also seen for salmon soup (data not shown).

**Table 5.** Debittering effect of whole proteome cutting in the cod proteome

| No. | Total protease mixture (g) | Bitterness (1hr) | Bitterness (6hr) | Bitterness (30hr) |
| --- | --- | --- | --- | --- |
| 1 | 0 | ++++ | ++++ | ++++ |
| 2 | 8.18 | +++ | ++ | ++ |
| 3 | 16.36 | +++ | + | + |
| 4 | 81.8 | +++ | + | + |

### Debittering effects of trout extract on cod bone soup

Freshly made 100ml cod bone soup was mixed with 100ml trout bone soup plus 200ml water. After boiled for 20 min and stayed for 1hour at the room temperature, the bitter taste was evaluated. It was found that the bitter taste was completely eliminated, with the evaluation score from (+++) or (++++) to (-). The experiment repeated three times and got the similar results.

### Debittering effects of trout extract on salmon bone soup

Freshly made 100ml salmon bone soup was mixed with 100ml trout bone soup plus 200ml water. After boiled for 20 min and stayed for 1hour at the room temperature, the bitter taste was evaluated. It was found that the bitter taste was also completely eliminated, with the evaluation score from (+++) or (++) to (-). The experiment repeated three times and got the similar results, and when the ratio of salmon bone soup to trout bone soup was 1:2, the debittering effect was getting slightly better than the 1:1 ratio.

### Debittering effects of trout extract on bitter melon soup

Fresh bitter mellon was mixed with water at 1:1 (w/w) and then smashed into soup. Take 100ml bitter melon soup and mix with 100ml trout bone soup plus 200ml water. After boiled for 20 min and stayed for 1hour at the room temperature, the bitter taste was evaluated. It was found that the bitter taste was also almost completely eliminated, with the score from (++++) to (+) or (-). The experiment repeated three times and got the similar results, and when the ratio of bitter melon soup to trout bone soup was 1:2, the debittering effect was getting slightly better than the 1:1 ratio.

## DISCUSSION

### Correlation between predicted bitter peptide contents and the real bitter taste

It is hard to evaluate the contribution of a single type of bitter peptide to the total taste of a food. Meanwhile, the total numbers and types of all potential bitter peptides cannot easily get associated with the food taste (Supplementary Table 1s and Fig1s; Fig.1). Maybe the bitter peptide density in a proteome (the bitter peptide total numbers divided by the size of the proteome) matters. However, the bitter peptide density of the six types of food all lies within the range 34.3∼43.8 per kb proteome, and apparently has no correlation with bitter taste levels. This is consistent with previous observations^7, 20^ that only those peptides with molecular weight less than 5000-6000 have a chance to be bitter. It is also a common sense that amino acids compositions of a food have no direct correlation with its bitter taste extent, though bitter mellon’s bitterness may be related with its relatively high ratios in several amino acids (Supplementary Fig2s).

**Figure 1.**
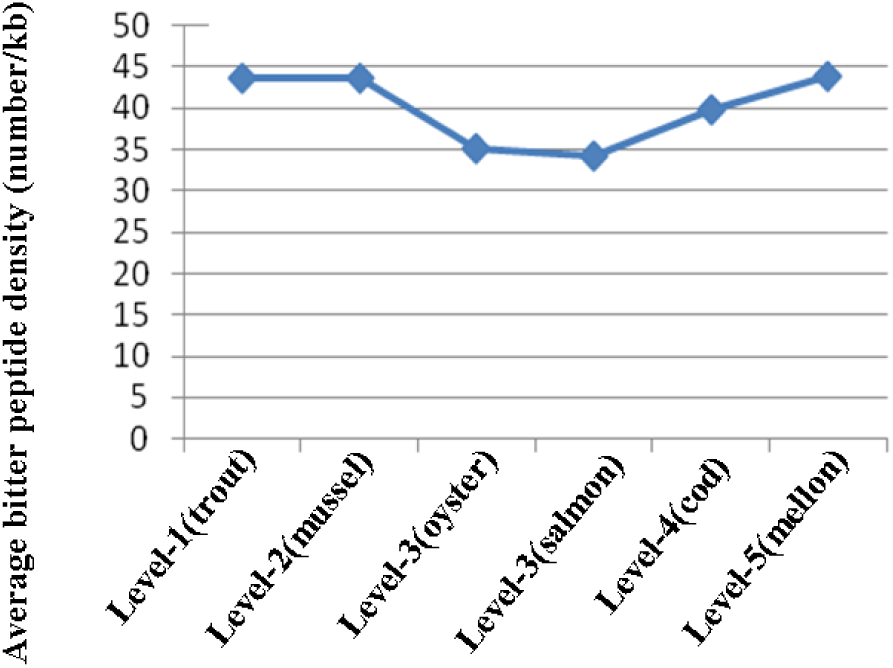
Predicted values of bitter peptide density for six different types of food that had big difference in the bitter taste. From level 1 to level 5, the bitterness increases from (-), (+), (++), (+++) to (++++). The mussel, oyster and mellon (bitter mellon) represented another three kinds of food locally collected.

### Which is better, digestion of the whole proteome or only bitter peptides?

Proteome-based strategy may be inefficient without careful control of protease digestion. One of the reasons is that some bitter peptides may become bitterer after broken into pieces; Some larger bitter peptides contain a more bitter small peptide within them. This is consistent with the observation that longer time (such as 24hr) of protease digestion of cod soup would bring more bitter taste than shorter time of digestion (such as 6hr). So, no matter whether the digestion is for the whole proteome or only for the bitter peptides, the residual bitterness can not be eliminated theoretically without facilitation of other ways, such as masking by trout soup. In practice, even the specifically designed bitter peptides-digestion regime still needs facilitation by optimization and control of digestion conditions to achieve satisfactory bitterness levels.

### Fatty acids may facilitate trout’s bitter-masking capacity

The bitter taste of cod fish was previously studied using conventional methods^21^ and the authors found that Flavourzyme^®^ did not reduce the bitterness well while the use of butanol and cholestyramine resin separately or in combination reduced the bitter taste from fish protein hydrolysates to levels hardly discernible in 1% concentration. In this study, the bitter taste of cod fish bone soup was masked completely by trout bone soup, suggesting that the latter has some substance to interact with bitter taste receptors or scavenging bitter substances themselves. There are several types of naturally occurred substances that bind with bitter taste receptors, such as some lipoproteins, cyclodextrin and cyclofructan.^22-24^ Among them, lipoproteins may be of a particular interest for this study because trout bone soup had a much higher lipid compositions as observed than other two kinds of bone soup, though the authors were still not sure whether lipoproteins^25^ or fatty acids^26^ (Table 6) played important roles for trout soup to be able of masking bitter taste. Table 6 clearly indicated that the trout bone soup had higher concentrations of about 20 different fatty acids than cod and salmon soups. According to several reports, some fatty acids can be used to efficiently mask bitter taste.^27-29^ Tomotake et al. (1998) and Fujita et al (2004) found that fatty acid salts such as sodium stearate, palmitate, and laurate in relatively high concentrations of about 1% were able to reduce the bitter taste of a 100-ppm quinine solution significantly, while there was 2.54% palmitic acids in the trout soup (Table 6) and may contribute to mask the bitter taste. Ryousuke Homma et al^30^ found that a 0.5mM mixture of palmitic acid, stearic acid and myristic acid significantly reduced the bitterness of quinine hydrochloride, while the concentration of the above mixture in the trout soup (99mM, 21mM and 28mM for the three types of fatty acids respectively) was about 40 times larger than that of the above mixture used for the sensory test, which shall be adequate to demonstrate the bitterness-masking effects. Though whether or how these fatty acids contribute to the bitterness-masking capacity of trout soup is still vague at this time, the literature strongly supports that fatty acids interwine with bitterness-sensing signaling pathway^31-34^ and likely play an important role for trout soup to mask the bitter taste.

**Table 6.** Fatty acid concentrations in three types of bone samples

|  | Fatty acid | Concentration (g kg<sup>-1</sup>) |  |  |
| --- | --- | --- | --- | --- |
|  |  | cod | trout | salmon |
| 1 | Heneicosanoic acid | ND | ND | ND |
| 2 | Tricosanoic acid | ND | ND | ND |
| 3 | trans-9-Octadecenoic acid | ND | ND | ND |
| 4 | trans-trans-9,12-Octadecadienoic acid | ND | ND | ND |
| 5 | Butyric acid | ND | ND | ND |
| 6 | Caproic acid | ND | ND | ND |
| 7 | Caprylic acid | ND | ND | ND |
| 8 | Capric acid | ND | ND | ND |
| 9 | Undecanoic acid | ND | ND | ND |
| 10 | Lauric acid | ND | ND | ND |
| 11 | Tridecanoic acid | ND | ND | ND |
| 12 | Myristic acid | 0.0028 | 0.064 | 0.017 |
| 13 | Pentadecanoic acid | ND | 0.0046 | ND |
| 14 | Palmitic acid | 0.015 | 0.254 | 0.034 |
| 15 | Margaric acid | ND | 0.0042 | ND |
| 16 | Stearic acid | 0.0026 | 0.06 | 0.0078 |
| 17 | Arachidic acid | ND | ND | ND |
| 18 | Behenic acid | ND | ND | ND |
| 19 | Lignoceric acid | ND | ND | ND |
| 20 | cis-9-Tetradecenoic acid | ND | ND | ND |
| 21 | cis-10-Pentadecenoic acid | ND | ND | ND |
| 22 | cis-9-Hexadecenoic acid | 0.0022 | 0.06 | 0.0142 |
| 23 | cis-10-Heptadecenoic acid | ND | ND | ND |
| 24 | cis-11-Octadecenoic acid | ND | 0.05 | 0.0056 |
| 25 | cis-9-Octadecenoic acid | 0.0146 | 0.668 | 0.034 |
| 26 | cis-11-Eicosenoic acid | 0.0046 | 0.007 | 0.022 |
| 27 | cis-13-Docosenoic acid | ND | 0.0084 | 0.0084 |
| 28 | cis-15-Tetracosenoic acid | ND | 0.0058 | 0.0022 |
| 29 | all cis-9,12,15-Octadecatrienoioc acid | ND | 0.0074 | ND |
| 30 | all cis-11,14-Eicosadienoic acid | ND | 0.004 | ND |
| 31 | all cis-11,14,17-Eicosatrienoic acid | ND | 0.042 | 0.024 |
| 32 | all cis-8,11,14,17-Eicosatetraenoic acid | ND | ND | ND |
| 33 | all cis-5,8,11,14,17-Eicosapentaenoic acid | ND | ND | 0.0036 |
| 34 | all cis-7,10,13,16,19-Docosapentaenoic acid | ND | 0.0068 | ND |
| 35 | all cis-4,7,10,13,16,19-Docosahexaenoic acid | ND | 0.0032 | 0.0022 |
| 36 | cis,cis-9,12-Octadecadienoic acid | ND | 0.062 | 0.0028 |
| 37 | all cis-6,9,12-Octadecatrienoic acid | ND | 0.0062 | ND |
| 38 | all cis-8,11,14-Eicosatrienoic acid | ND | 0.003 | ND |
| 39 | all cis-5,8,11,14-Eicosatetraenoic acid | ND | ND | ND |
| 40 | all cis-13,16-Docosadienoic acid | ND | 0.0024 | ND |
| 41 | Total fat | 0.06 | 1.54 | 0.22 |

Note: ND: not detected

## CONCLUSION

In this study, 46 known bitter peptides in the cod proteome were targeted for specific protease digestion to eliminate bitter taste from the cod bone soup. Though the debittering effect was apparent, the bitter taste was not completely removed. However, the total bitter taste can be surprisingly removed simply by addition of trout extract. The strong debittering power of trout extract was further confirmed by the debittering tests on salmon bone soup and bitter melon, both with perfect results.^35^ These results indicated that the cod bone soup bitterness comes not only from bitter peptide but also from other substances that can be masked by trout extract. Considering the fact that trout bone soup had much higher lipid content than cod and salmon bone soups, fatty acids are probable substances that mediate the bitterness-masking and remain worthy to be determined in the future.

## Supplementary materials

**Supplementary Fig1s.**
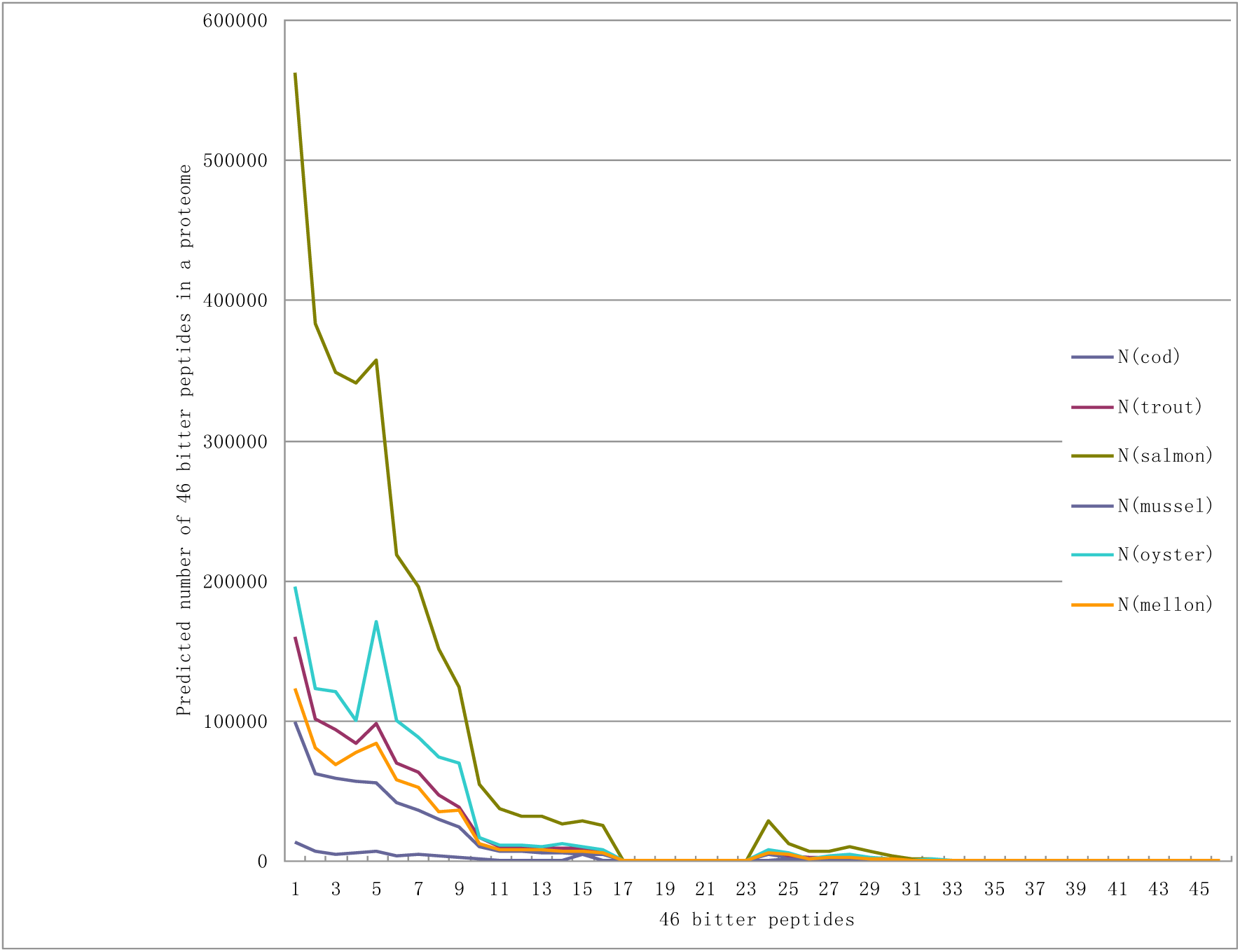
Predicted numbers of 46 bitter peptides in the six proteomes. The sequences of 46 bitter peptides can be found in Table 1.

**Supplementary Fig2s.**
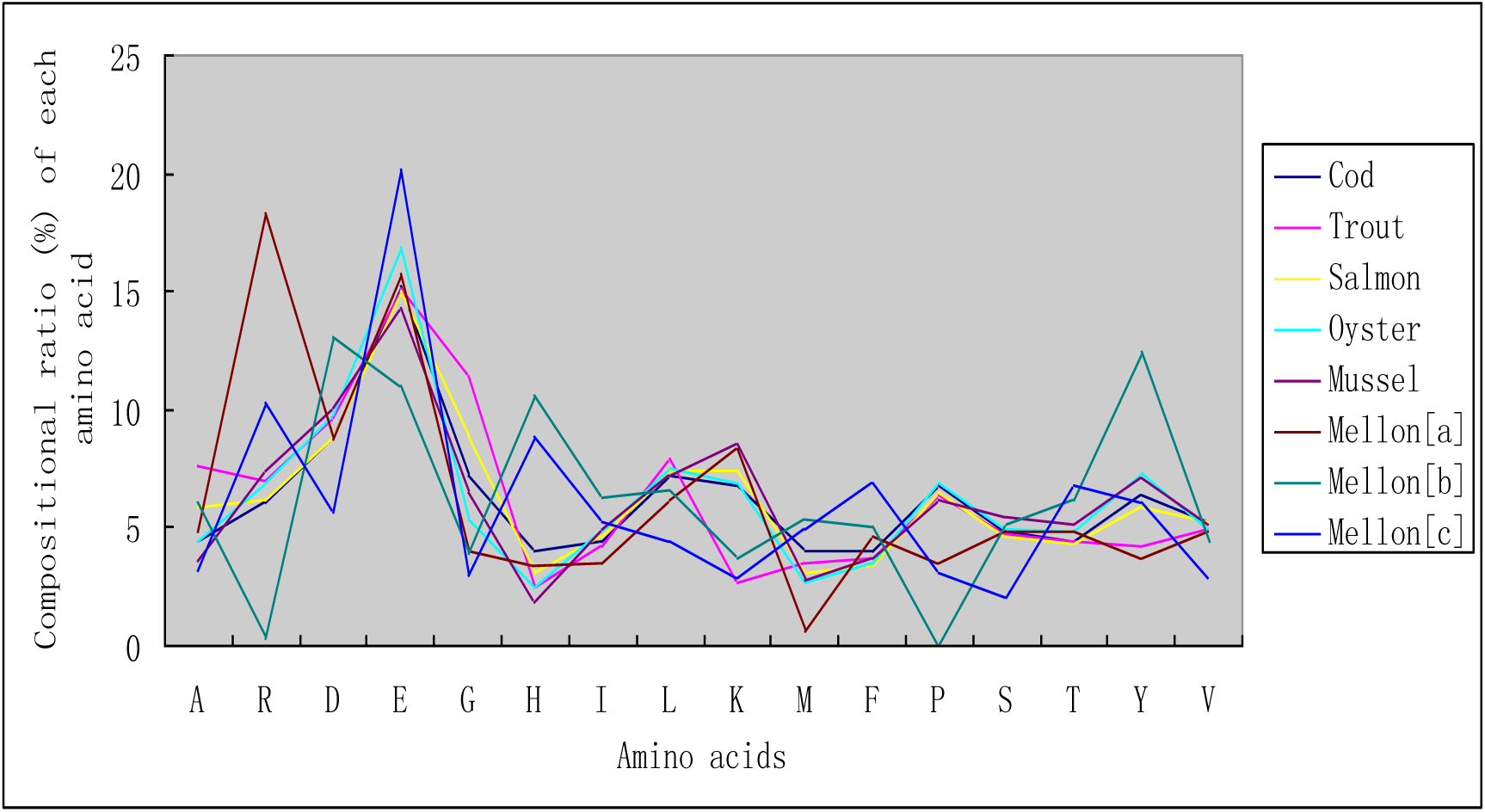
Amino acids compositions (percentage of each amino acid to the whole amino acids) of six types of food. Cod, trout and salmon represented three different bone samples. Oyster and mussel represented the whole body samples except for the shells. The values were the average of three repetitions for the above five types of sample. Mellon [a,b,c] represented three cases of amino acids concentration measurement of bitter mellon as the following literature: [a] Effects of heat drying process on amino acid content of Momordica charantia L. *Amino Acids& BioticResource*.2010, 32(2):14-16.[b] Quantitative analysis of protein, amino acids and inorganic elements in Momordicacharantia. *Bullet. Hunan Univ.* 1996,21(5): 305-307.[c] Effect of microwave treatment on quality of Momordica charantia L. *Mod. Food Sci. Technol.*2009, 25(11): 1250-1253.

**Supplementary Table 1s.**
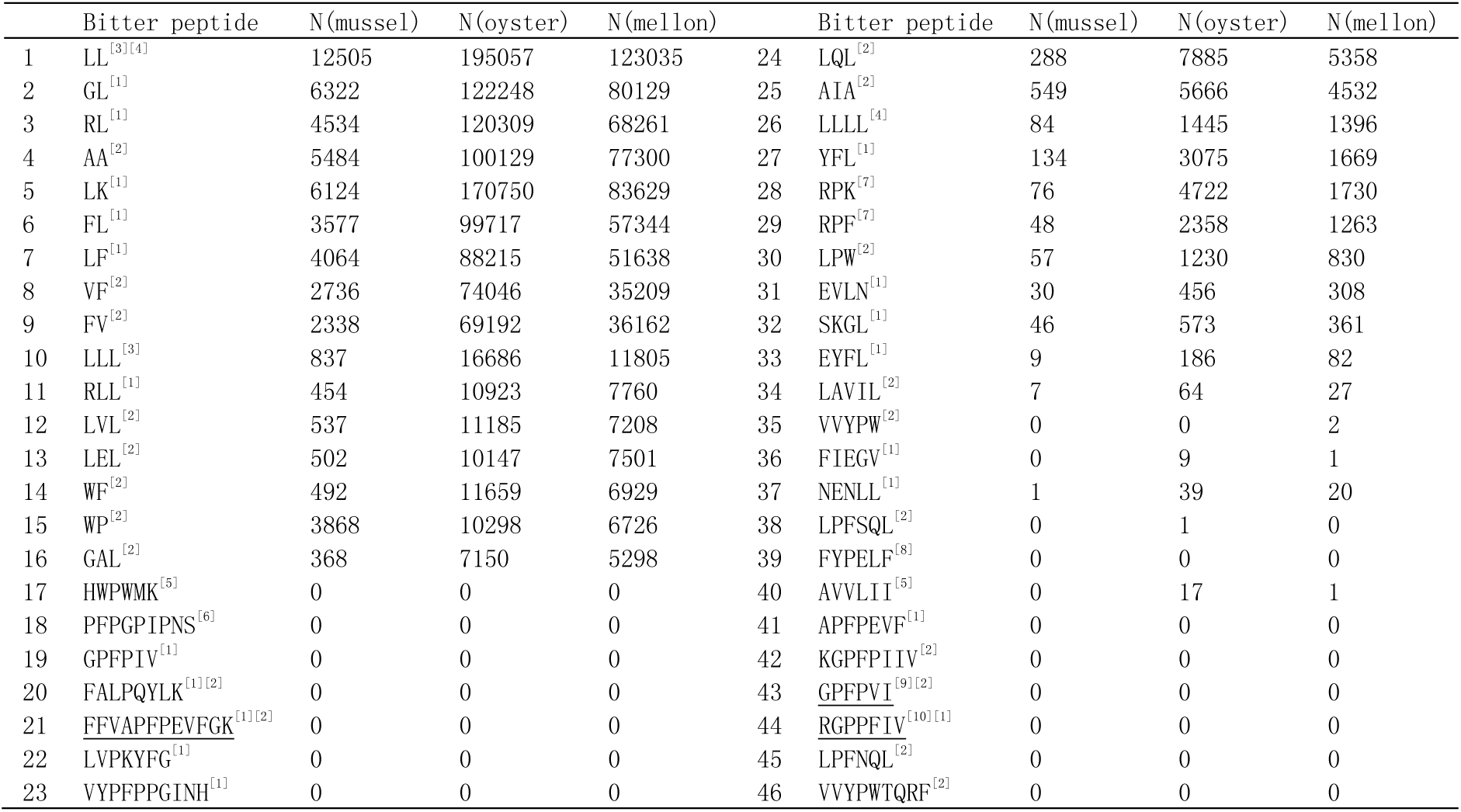
Predicted numbers of 46 bitter peptides in mussel, oyster and bitter mellon in their proteomes

